# Neurocognitive Heterogeneity in Social Anxiety Disorder: The Role of Self-Referential Processing and Childhood Maltreatment

**DOI:** 10.1101/2020.09.22.307835

**Authors:** Anat Talmon, Matthew Luke Dixon, Philippe R. Goldin, Richard G. Heimberg, James J. Gross

**Affiliations:** Stanford University, CA, USA; University of California Davis CA, USA; Temple University, Philadelphia, PA, USA

## Abstract

Social anxiety disorder (SAD) is characterized by negative self-beliefs and altered brain activation in the default mode network (DMN). However, the extent to which there is neurocognitive heterogeneity in SAD remains unclear. We had two independent samples of patients perform a self-referential encoding task, and complete self-reports of childhood maltreatment, subjective well-being, and emotion regulation. In the replication sample, we also measured DMN activation using functional magnetic resonance imaging. K-means clustering revealed two distinct sub-groups of SAD patients in the discovery sample. Cluster 1 demonstrated higher levels of negative and lower levels of positive self-referential trait endorsement, and significantly higher levels of childhood emotional maltreatment, lower subjective well-being, and altered emotion regulation strategy use. A similar pattern was observed in the replication sample, which further demonstrated higher DMN activation during negative trait judgments in cluster 1. These findings reveal neurocognitive heterogeneity in SAD and its relationship to emotional maltreatment.

## INTRODUCTION

Substantial heterogeneity exists within patient populations that receive a particular clinical diagnosis. This heterogeneity may reflect variation in etiology and have implications for current well-being, prognosis, and treatment. This has fueled interest in defining neurocognitive subtypes that capture some of this heterogeneity. Studies have increasing turned to unsupervised (data-driven) clustering approaches to identify potential sub-groups within particular disorders (Feczko et al., 2019; Kaczkurkin et al., 2020; Marquand, Wolfers, Mennes, Buitelaar, & Beckmann, 2016).

One potentially promising target is social anxiety disorder (SAD). SAD is the most common anxiety disorder, with a 12.1% lifetime prevalence (Stein & Stein, 2008), and it is characterized by an intense, persistent fear of being evaluated in social situations (Heimberg et al., 2014). Heterogeneity within SAD patients has not been explored in detail, although it was suggested that patients may vary in the types of self-related thoughts and beliefs they exhibit (Gregory & Peters, 2017), and in self-definition, with a specific emphasis on self-criticism and dependency (Kopala-Sibley, Zuroff, Russell, & Moskowitz, 2014). Low remission rates following therapy (Steinert, Hofmann, Leichsenring, & Kruse, 2013) also point to possible heterogeneity.

One of the core attributes of this disorder is negative self-beliefs (Clark & Wells, 1995; Hofmann, 2007; Moscovitch, 2009; Rapee & Heimberg, 1997). While healthy controls generally show a positivity bias, SAD patients generally show a negativity bias (P. R. Goldin, Manber-Ball, Werner, Heimberg, & Gross, 2009). Indeed, previous studies demonstrated that negative self-referential processing is a major component of patients’ phenotype (Abraham et al., 2013; Button, Browning, Munafò, & Lewis, 2012; M. L. Dixon et al., 2020;P. Goldin, Ramel, & Gross, 2009; P. Goldin, Ziv, Jazaieri, & Gross, 2012) and has implications for treatment response (Thurston, Goldin, Heimberg, & Gross, 2017). Self-referential processing is commonly associated with engagement of the default mode network (DMN) (Andrews-Hanna, Smallwood, & Spreng, 2014; Buckner, Andrews-Hanna, & Schacter, 2008; Northoff et al., 2006), a system that shows aberrant activation patterns in SAD (Bruehl, Delsignore, Komossa, & Weidt, 2014; M. L. Dixon et al., 2020). While SAD patients may show excessive negative self-beliefs and altered DMN recruitment on average, we hypothesized that there might be important individual variability, with some patients exhibiting more extreme negative bias than others.

If it is possible to identify neurocognitive sub-groups of SAD patients, a critical question concerns the possible relationship between such groups and life events. One possibility is that clinical heterogeneity within SAD may relate to differences across the lifespan in exposure to traumatic events, such as childhood maltreatment. Prior work has demonstrated that childhood maltreatment is associated with anxiety and depression (Iffland, Sansen, Catani, & Neuner, 2012; Simon et al., 2009; Vachon, Krueger, Rogosch, & Cicchetti, 2015), later identity diffusion (i.e., when a person’s identity remains unresolved and not fully formed), altered self-perception (Scott et al., 2014), and negative self-referential processing (Penner, Gambin, & Sharp, 2019; Toth, Cicchetti, MacFie, Maughan, & Vanmeenen, 2000). Given the known relationships between childhood maltreatment, psychopathology, and altered self-referential processing, we hypothesized that heterogeneity in SAD, in the form of more extreme negative self-beliefs and altered DMN activation, might be associated with greater exposure to childhood maltreatment.

In the current study, we combined theoretically-informed hypotheses with an unsupervised clustering approach to investigate potential heterogeneity within SAD. We used large discovery (N = 95) and replication (N = 97) samples from independent sets of patients to validate our findings. We first clustered patients based on behavioral data in a self-referential task, and then subsequently clustered them based on behavioral and brain data (default mode network activation) to assess the added value of neuroimaging data beyond behavioral data alone. After identifying two clusters (sub-groups), we investigated the potential role of childhood maltreatment in differentiating the sub-groups and compared sub-groups on measures of general well-being and adaptive functioning.

## Materials and methods

### Participants

Two independent samples of SAD patients were included in data analysis (M. L. Dixon et al., 2020; P. Goldin, W. Ramel, et al., 2009; P. R. Goldin et al., 2009). The discovery and replication samples included 95 patients with SAD (mean [SD] age = 33.53 [8.56] years; 46 (48.4%) women) and 97 patients with SAD (mean [SD] age = 32.94 [8.08] years; 51 (53.1%) women), respectively. Patients provided informed consent in accordance with the Institutional Review Board at Stanford University, passed MRI safety screening, were 22 to 55 years of age, were fluent in English, and were right-handed. Patients with SAD met criteria for a primary diagnosis of generalized SAD based on the Anxiety Disorders Interview Schedule for DSM-IV: Lifetime version (Di Nardo, Brown, & Barlow, 1994) (for additional details and screening criteria see methods in the **Supplementary Information, SI**). Severity of social anxiety and avoidance was measured with the Liebowitz Social Anxiety Scale Self-Report (LSAS-SR) (Liebowitz & Pharmacopsychiatry, 1987).

### Self-Referential Encoding Task

Participants performed a self-referential encoding task (Derry & Kuiper, 1981) (see **SI** for details). Stimuli were 25 positive and 25 negative social trait adjectives from the Affective Norms for English Words database (Bradley & Lang, 1999). Participants viewed the trait words and made a yes/no judgment indicating whether the trait was self-descriptive (‘self’ judgment condition), or a yes/no judgment indicating whether the trait was written in all upper-case letters (‘case’ judgement condition). There were five blocks of each of the four trial types. Each block started with a question screen (either “Describes ME?” or “UPPER case?”) for 1.5 seconds, and then five positive or five negative adjectives were presented one at a time for 3 seconds each. Participants made a yes/no response using a button pad during presentation of each of the five stimuli. In total, participants made 25 judgments for each of the four trial types. At the end of the run there was a 3-second fixation cross and a 3-second blank screen. Stimulus order was pseudo-randomized in terms of block sequence, with no more than two blocks of the same condition in a row. The sequence of words as well as whether they were upper or lower case were randomized within each block. For each participant we calculated the percentage of positive and negative trait words that were endorsed during the self judgment conditions and mean accuracy during the case judgment conditions.

Only behavioral data were collected for the discovery sample. Behavioral and functional magnetic resonance imaging (fMRI) data were collected for the replication sample. Only participants who scored 70% correct or higher on the case judgement trials were included in further analyses to ensure that they understood the task and were paying attention. This criterion led to the exclusion of 9 patients in the discovery sample (8.7% of sample) and 17 patients in the replication sample (14.9% of sample), resulting in final samples of N = 95 and N = 97, respectively.

### Measure of Childhood Maltreatment

Childhood maltreatment was assessed with the Childhood Trauma Questionnaire(Bernstein et al., 2003), which consists of 28 items reflecting five forms of childhood maltreatment: physical abuse, sexual abuse, emotional abuse, physical neglect, and emotional neglect. Ratings were made on a 5-point scale ranging from 1 (*never true*) to 5 (*very often true*). Sum scores were used, with higher scores representing greater levels of childhood maltreatment. Internal consistencies of this scale were adequate (Bernstein & Fink, 1998; Bernstein et al., 2003). Since emotional abuse and emotional neglect were highly correlated (*r* = .69), we calculated a mean score representing emotional trauma. We did the same for physical abuse and neglect (*r* = .49).

### Measures of Well-being and Adaptive Functioning

*The Satisfaction with Life Scale*(Diener, Emmons, Larsen, & Griffin, 1985) consists of five items that assess a person’s satisfaction with life in general (e.g., “In most ways my life is close to ideal”), rated on a 7-point Likert-type scale ranging from 1 (*very much opposed*) to 7 (*strongly agree*). Reported internal consistency and two month test-retest reliability for scores on the SWLS were .87 and .82, respectively (Diener et al., 1985).

*The Perceived Stress Scale* (Cohen, Kamarck, & Mermelstein, 1983) consists of 14 items that assess the degree to which situations in a person’s life are appraised as stressful (e.g., “In the past month, how often have you felt nervous and “stressed out”?”), rated on a 5-point Likert-type scale ranging from 1 (*never*) to 5 (*very often*). Internal consistency and construct validity of this scale were previously supported (Roberti, Harrington, & Storch, 2006).

*The Emotion Regulation Questionnaire* (J. J. Gross & O. P. John, 2003) consists of 10-items that assess both suppression (e.g., “I keep my emotions to myself”) and reappraisal (e.g., “When I want to feel less negative emotion (such as sadness or anger), I change what I’m thinking about”) strategies, rated on a 7-point Likert-type scale ranging from 1 (*strongly disagree*) to 7 (*strongly agree*) based on frequency of using each strategy. Convergent and discriminant validity of this scale were previously supported (J. J. Gross & O. P. John, 2003).

*The Emotion Regulation Questionnaire-Self Efficacy* (P. Goldin, Manber, Hakimi, Canli, & Gross, 2009) assesses how capable participants believe they are of using reappraisal and/or suppression when they really want to, using the same item set described above. Participants rated their agreement or disagreement with each item on a scale from 1 (strongly disagree) to 7 (strongly agree). This scale has good reliability and construct validity (J. Gross & O. John, 2003).

### fMRI Data Analysis

#### Acquisition

In the replication sample only, fMRI data were collected using a General Electric 3T Signa magnet with a T2*-weighted gradient-echo spiral-in/out pulse sequence (Glover & Law, 2001). Twenty-four ascending interleaved axial slices were acquired (4.5 mm slice thickness; single shot; repetition time (TR) = 1.5s; echo time (TE) = 28.5ms; flip angle (FA) = 65°; field of view (FOV) = 220 mm; matrix size = 64 × 64, voxel resolution 3.438 mm2 x 4.5 mm). Each patient completed one functional run during which 230 functional volumes were acquired. Data collected during the first 4 TRs were discarded to allow for equilibration effects. Before functional imaging, a high resolution T1-weighted structural image was acquired using fast spin-echo spoiled gradient recall (132 slices; TR = 3s; TE = 68ms; FOV = 220mm; matrix size: 256 x 256; voxel size = 1 x 1 x 1.2mm). Head movement was restricted using a bite-bar and foam padding.

#### Preprocessing

Using SPM12, the data were corrected for motion via realignment to the first volume (using a 6-parameter rigid body transformation), and slice-time corrected (to the middle slice). Each subject’s T1 image was bias-corrected and segmented using a nonlinear deformation field to map it onto template (ICBM) tissue probability maps for gray/white matter and CSF. Parameters obtained from this step were subsequently applied to the functional data (re-sampled to 3 mm3 voxels) during normalization to MNI space. The data were spatially-smoothed using an 8-mm3 full-width at half-maximum Gaussian kernel to reduce the impact of inter-subject variability in brain anatomy.

#### First-level analysis

Multiple regression analyses were conducted at the first-level using the following regressors convolved with a canonical hemodynamic response function: (i) instruction (question) cue; (ii) negative self judgment; (iii) positive self judgment; (iv) negative case judgment; (v) positive case judgment. To account for residual noise, the model also included 6 motion parameters from realignment and framewise displacement timecourse. The model included constants to account for between-run differences in mean activation and a high-pass filter (128s cutoff) to remove low-frequency drifts.

We focused on DMN activation given its theoretical relevance to self-referential processing and SAD. We extracted and averaged mean beta values from two regions identified as core nodes of the DMN (Andrews-Hanna, Reidler, Sepulcre, Poulin, & Buckner, 2010) using coordinates from a meta-analysis of self-referential processing (Northoff et al., 2006). These regions were the medial prefrontal cortex (10mm sphere centered on the coordinates: x = −2, y = 49, z = 7; and posterior cingulate cortex (10mm sphere centred on the coordinates: x = −3, y = −61, z = 31 (**Figure S1**). We extracted mean beta values separately for the positive self > case judgment and negative self > case judgment contrasts.

### Clustering Analysis

We used k-means clustering, implemented in Matlab (R2017B), to identify sub-groups of SAD patients. Given our sample size, and the fact that specifying a higher number of clusters would have resulted in clusters with low participant numbers and would run the risk of overfitting, we restricted the analysis to a 2-cluster solution. The initial input features to the clustering algorithm were the percentage of positive and negative trait words endorsed by each participant. Subsequently, we added mean activation within the default mode network during positive and negative judgements as additional input features. To ensure that all input features had equal weight, data were Z-scored prior to clustering. By using a small number of input features and testing the clustering approach in independent samples, we maximized the likelihood of discerning replicable sub-groups.

### Statistical Analyses

The two clusters were compared on relevant outcome variables using independent samples t-tests using two-tailed p-values. For the discovery sample, p-values for the comparison of clusters in relation to childhood maltreatment were Bonferroni corrected for the three tests performed. P-values for the comparison of clusters in relation to well-being measures were corrected based on the false discovery rate (FDR). Replication findings were considered significant at *p* = .05 (two-tailed).

## RESULTS

### Preliminary Analyses

A description of demographics for the two samples, as well as correlations among study variables, are presented in Tables 1 and 2 in the **SI**.

### Discovery Sample

SAD patients were divided into two clusters based on their profile of SRET trait endorsement scores. 55.8% (*N* = 53) of participants were classified as belonging to cluster 1 and 44.2% (*N* = 42) were classified as belonging to cluster 2 (**Figure 1A**). Participants in cluster 1 (negative self) compared to those in cluster 2 (positive self) endorsed fewer positive traits (M_Cluster 1_ = 24.62%; M_Cluster 2_ = 61.33%) and more negative traits (M_Cluster 1_ = 69.53%; M_Cluster 2_ = 34.34%). This clustering pattern reveals substantial heterogeneity within the SAD sample, with sub-groups that demonstrate marked differences in self-beliefs. Although SAD is thought to be associated with primarily negative self-beliefs, here we discover a sub-group with overall positive self-beliefs. Notably, the two clusters did not differ on demographic characteristics [years of education: *t*(89) = −1.61, *p* = .11; age: *t*(93) = −.65, *p* = .52; sex: *X*^2^ (1, N = 95) = .02, *p* = .89].

**Figure 1.**
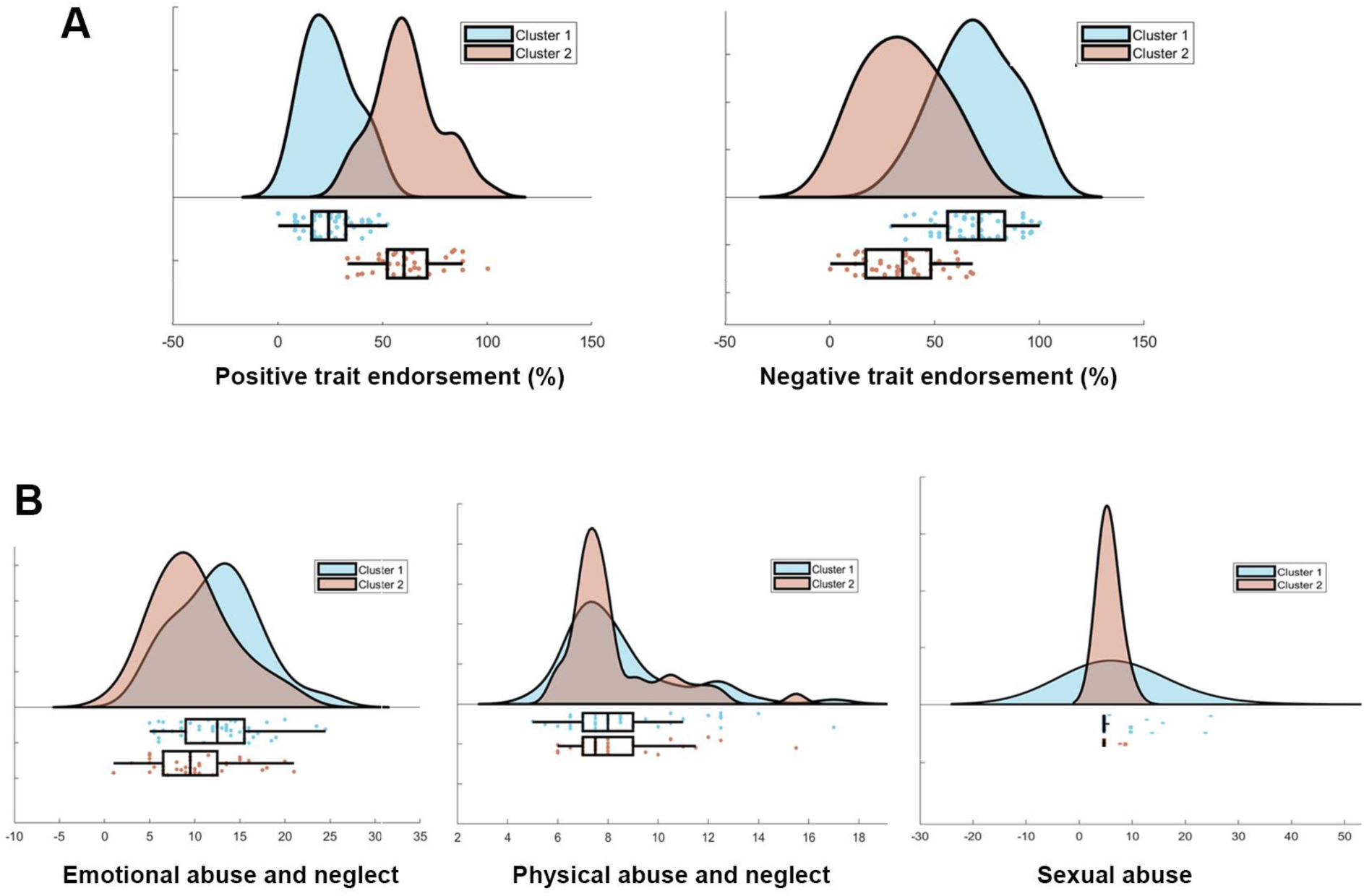
Discovery sample SAD clusters show distinct profiles of positive and negative self-referential trait endorsement. (A) Cluster 1 compared to cluster 2 patients were less likely to endorse positive traits and more likely to endorse negative traits. (B) The clusters also differed in exposure to emotional abuse and neglect, but not physical abuse and neglect or sexual abuse. Raincloud plots were generated using open source code(Allen, Poggiali, Whitaker, Marshall, & Kievit, 2019).

We next compared the clusters with respect to self-reported levels of exposure to three types of childhood maltreatment (**Figure 1B**). These analyses revealed that the two clusters differed in self-reported emotional maltreatment [*t*(90) = 2.81, *p* = .02, Bonferroni corrected], with cluster 1 (negative self) reporting higher levels of emotional maltreatment (*M* = 12.54, *SD* = 4.57) than cluster 2 (positive self) (*M* = 9.87, *SD* = 4.50). Breaking down emotional maltreatment, we found that the clusters differed in both emotional abuse [*t*(90) = 2.01, *p* = .048] and emotional neglect [*t*(86) = 2.54, *p* = .01]. Notably, the two clusters did not differ in levels of self-reported exposure to physical maltreatment [*t*(90) = .59, *p* = 1.67, Bonferroni corrected], or exposure to sexual abuse [*t*(90) = 1.77, *p* = .25, Bonferroni corrected]. This reveals a potentially selective relationship between the SAD groups we identified and early-life exposure to emotional trauma.

The clusters differed in self-reported social anxiety severity (**Figure S2A**). Cluster 1 (negative self) reported greater severity (M = 89.16, *SD* = 20.46) than cluster 2 (positive self) (M = 78.76, *SD* = 16.29) [*t*(92) = 2.68, *p* = .01].

To illuminate the functional significance of the identified clusters, we performed exploratory analyses comparing the clusters on subjective well-being and emotion regulation (**Figure 2**). Cluster 1 (negative self) compared with cluster 2 (positive self) reported lower satisfaction with life [*t*(92) = 5.85, *p* < .001, FDR corrected], and higher stress [*t*(92) = 3.75, *p* < .001, FDR corrected]. Cluster 2 (positive self) reported more use of reappraisal to regulate emotion in daily life than cluster 1 (negative self) [*t*(85) = 2.99, *p* < .001, FDR corrected]. The clusters did not differ in perceived self-efficacy of using reappraisal [*t*(29) = 1.01, *p* = .39], or the frequency [*t*(85) = 1.05, *p* = .39, FDR corrected] or self-efficacy [*t*(85) = .09, *p* = .93, FDR corrected] of using suppression as an emotion regulation strategy. These results reveal that the clusters defined based on self-referential processing also differ in some aspects of subjective well-being and emotion regulation strategy use.

**Figure 2.**
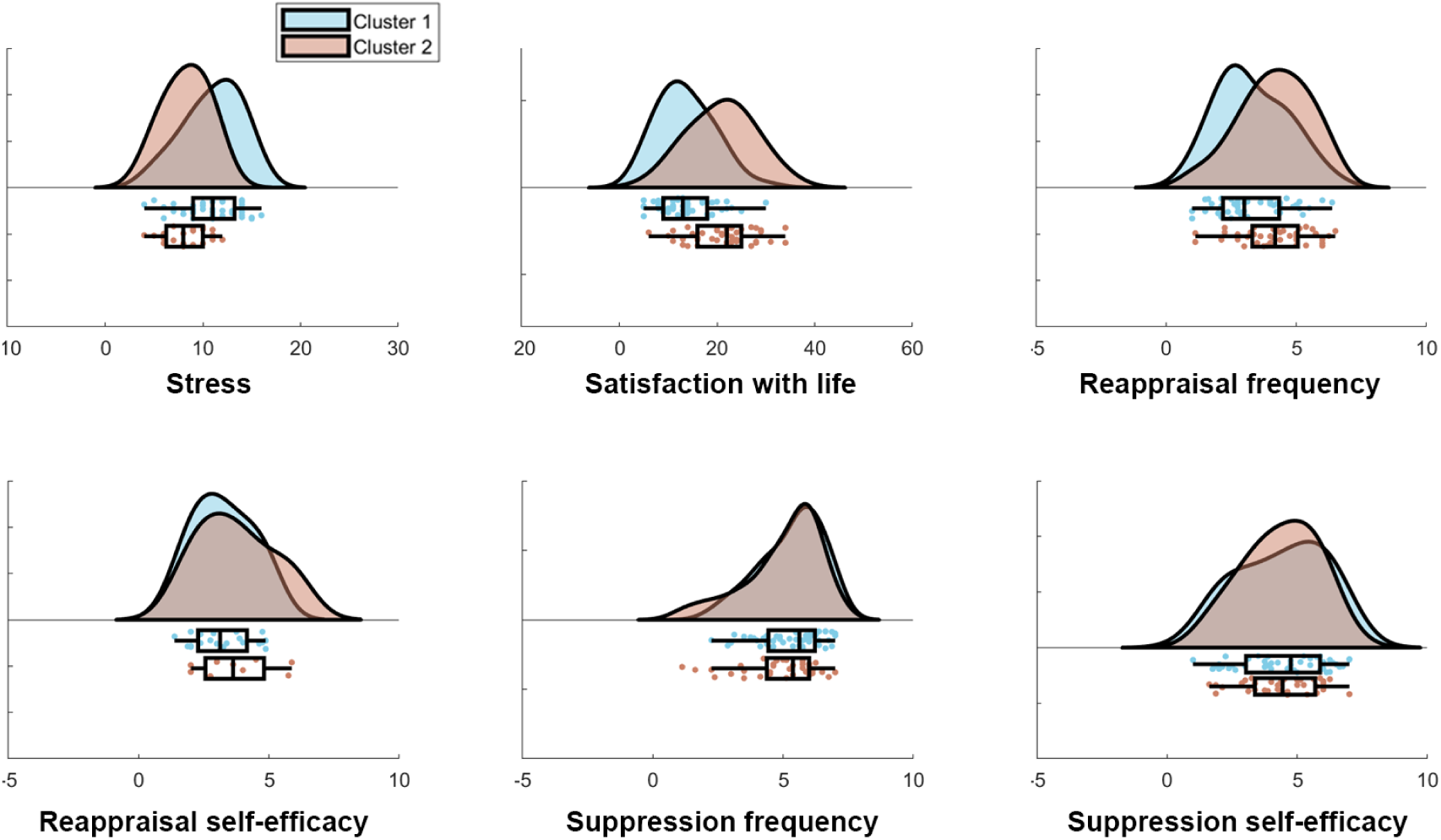
Subjective well-being and emotion regulation strategy use as a function of SAD cluster in the discovery sample.

### Replication Sample

Results from the replication sample supported the major findings from the discovery sample. In this case, 73.2% (*N* = 71) of participants were classified as belonging to cluster 1 (negative self) and 26.8% (*N* = 26) were classified as belonging to cluster 2 (positive self), again revealing sub-groups with marked differences in self-beliefs (**Figure S3A**). Replicating the discovery sample results, the clusters differed in emotional maltreatment [t(95) = −2.24, *p* = .03], but not physical maltreatment [t(95) = .90, *p* = .40], or sexual abuse [t(95) = .18, *p* = .45] (**Figure S3B**).

There was also a replication of group differences in well-being variables (**Figure S4**), with cluster 1 (negative self) compared to cluster 2 (positive self) reporting lower satisfaction with life [*t*(95) = 6.27, *p* < .001] and more stress [*t*(95) = −3.88, *p* < .001]. There was no difference between clusters in the frequency or self-efficacy of using reappraisal or suppression (all *p*’s > .2). There was no difference between clusters in social anxiety severity (**Figure S2B)** [*t*(95) = −.95, *p* = .34] or demographic characteristics [years of education: *t*(93) = .11, *p* = .91; age: *t*(94) = .92, *p* = .36; gender: *X*^2^ (1, N = 96) = .11, *p* = .74].

### Replication Sample with Brain Data

Clustering participants based on behavioral and brain data (DMN activation) resulted in slightly more balanced sub-groups (in terms of sample size) than without the inclusion of brain data, with 63.8% (*N* = 60) of participants belonging to cluster 1 (negative self) and 36.2% (*N* = 34) belonging to cluster 2 (positive self) (**Figure 3A**). Beyond the differences in trait endorsement, cluster 1 compared to cluster 2 exhibited higher DMN activation during negative trait judgments (**Figure 3B**). The two clusters differed in self-reported emotional maltreatment [*t*(92) = −2.66, *p* = .01], but not physical maltreatment [*t*(92) = −.45, *p* = .75], or sexual abuse [*t*(92) = .28, *p* = .78] (**Figure 3C**).

**Figure 3.**
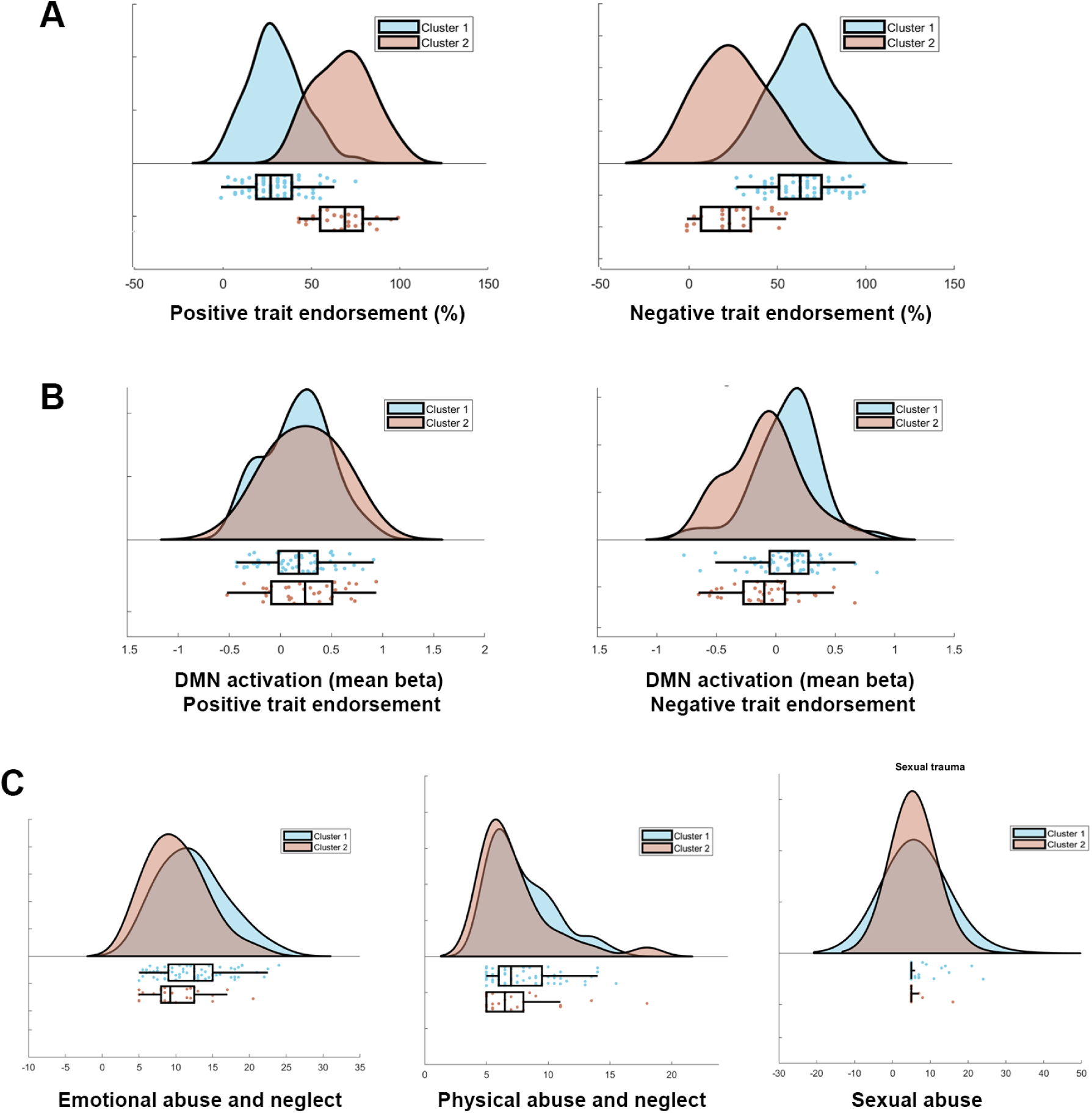
Replication sample SAD clusters show distinct profiles of positive and negative self-referential trait endorsement. (A) Cluster 1 compared to cluster 2 patients were less likely to endorse positive traits and more likely to endorse negative traits. (B) Cluster 1 compared to cluster 2 patients demonstrate greater default mode network (DMN) activation during negative trait judgments. (C) The clusters also differed in exposure to emotional abuse and neglect, but not physical abuse and neglect or sexual abuse.

The pattern of findings for the well-being measures was similar to those previously reported (**Figure 4**), however, in this case, the clusters did significantly differ in the frequency of using reappraisal [*t*(92) = 2.79, *p* = .01] (similar to the discovery sample). The clusters did not differ in social anxiety severity (**Figure S2C**) [*t*(92) = −.93, *p* = .36] or demographic characteristic [years of education: *t*(90) = 1.41, *p* = .16; age: *t*(90) = .83, *p* = .41; gender: *X*^2^ (1, N = 92) = 1.02, *p* = .31].

**Figure 4.**
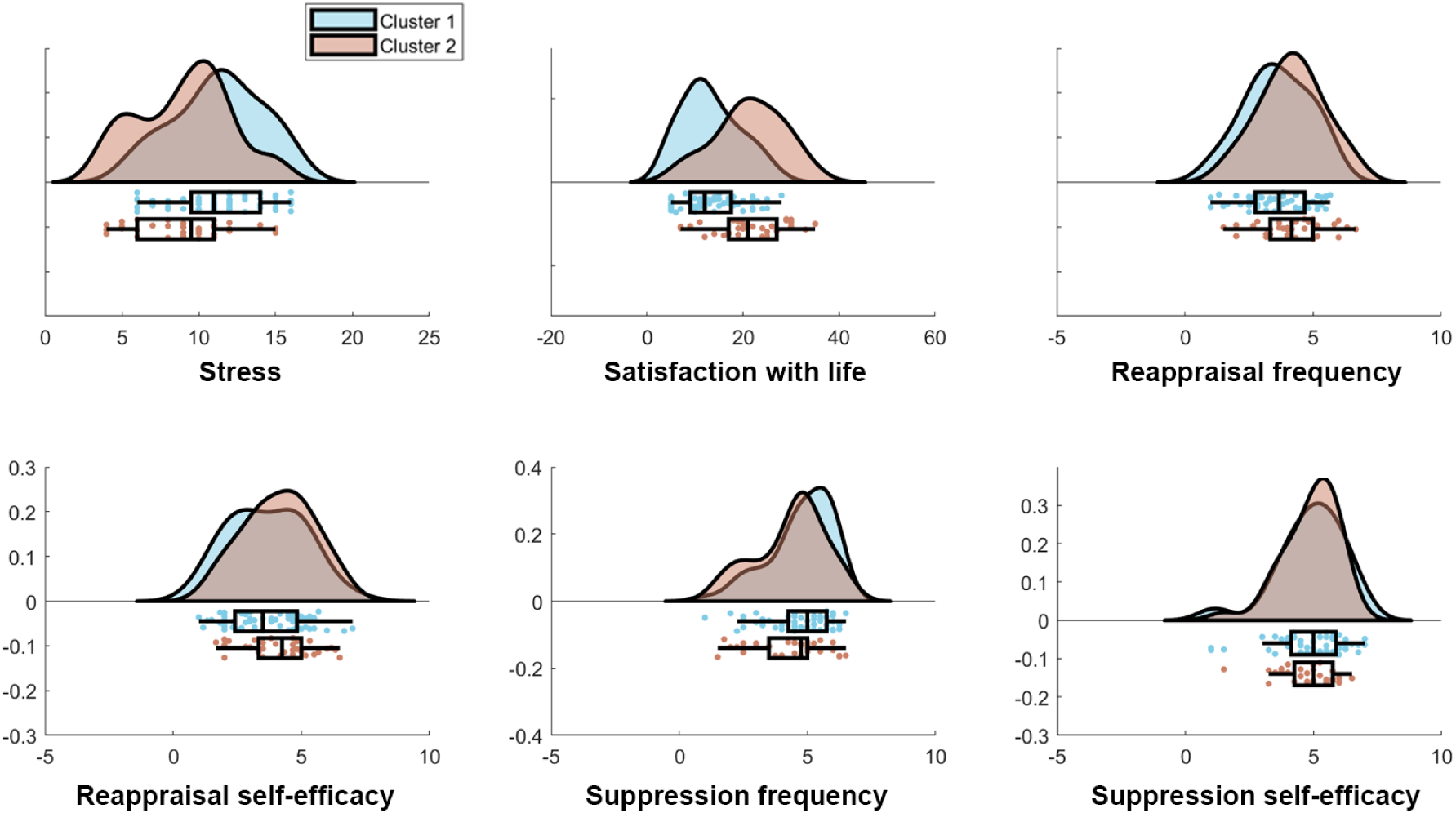
Subjective well-being and emotion regulation strategy use as a function of SAD cluster in the replication sample.

## DISCUSSION

The current study combined a data-driven approach with a constrained set of theoretically-relevant input features (positive and negative self-beliefs and default mode network activation) to reveal significant heterogeneity within a population of clinically diagnosed SAD patients. Although SAD patients demonstrate predominantly negative self-beliefs on average (P. R. Goldin et al., 2009), we identified two distinct clusters of patients; one of which demonstrated the expected pattern of predominantly negative self-beliefs, while the other demonstrated predominantly positive self-beliefs. The clusters also differed in default mode network activation strength during negative self-trait judgments.

These clusters significantly differed in level of exposure to childhood emotional abuse and neglect, but did not differ in level of exposure to physical or sexual trauma. This reveals a selective relationship between SAD sub-groups and emotional trauma and suggests that early life experiences may play a role in driving heterogeneity within SAD. We also found that the clusters differed in general well-being and adaptive functioning. The cluster demonstrating predominantly negative self-views showed lower satisfaction with life, higher stress, and less frequent use of an adaptive emotion regulation strategy (reappraisal) in daily life. We found that inclusion of neuroimaging data resulted in more balanced and differentiated clusters, suggesting that a combination of behavioral and brain data may provide the most insight into heterogeneous SAD sub-groups.

In the discovery sample – but not the replication sample – we found that the clusters differed in the severity of social anxiety symptoms. It is therefore unclear whether the clusters reflect overall symptom burden, or whether they are specifically related to self-referential processing in particular (distinct from other symptoms). Regardless, these data clearly reveal that self-beliefs and default mode network activation may be key elements underlying neurocognitive heterogeneity. Self-referential processing is known to affect attention (Zhao, Uono, Yoshimura, & Toichi, 2015), valuation (Berkman, Livingston, & Kahn, 2017; M. Dixon, Thiruchselvam, Todd, & Christoff, 2017), and decision-making (Johnson et al., 2005; Sui & Humphreys, 2015) and may therefore serve as a critical process that mediates interactions between the individual and their environment. The extent to which negative versus positive traits are incorporated into one’s self-concept may reflect one’s personal history of interaction with others. Childhood emotional trauma may significantly impact an individual’s emerging self-concept (Davis, Petretic-Jackson, & Ting, 2001; Frewen et al., 2011) and may bias learning such that negative feedback from others becomes more salient than positive feedback – a pattern observed in SAD (Koban et al., 2017). This may create a perpetuating effect, leading to a predominantly negative self-concept.

Several aspects of our study design are noteworthy. First, we took advantage of the simplicity of probing self-referential processing using a widely used task that is easy to administer. This may provide an alternative to DSM-defined symptoms for grouping patients into meaningful sub-groups. Moreover, given that negative self-beliefs are common to other disorders (e.g., depression) (Nejad, Fossati, & Lemogne, 2013), it may provide a transdiagnostic feature for discerning neurocognitive sub-types. Second, we used two large samples of independently acquired SAD patients and used a theoretically-informed clustering approach based on a small number of input features. This methodology allowed us to identify easily interpretable and generalizable clustering patterns.

### Limitations

Several limitations should be noted. First, we divided SAD patients into clusters based on a simple set of features. While this has noted advantages, there may be a multitude of features that may be informative about SAD sub-groups and could potentially provide even more detail about how such heterogeneity relates to measures of well-being and treatment response. Second, we used a limited number of well-being outcome variables. Future research could investigate other domains of well-being to provide a more comprehensive picture. Third, additional insight could be gleaned by including patients with a variety of disorders to look for transdiagnostic sub-groups that differ in self-referential processing. Finally, the use of a cross-sectional rather than longitudinal design and retrospective measure of childhood maltreatment precludes causal interpretations about its relationship with SAD heterogeneity.

## Acknowledgements

This research was supported by National Institute of Mental Health Grants R01 MH092416 and R01 MH76074 awarded to James Gross.

