## Supplementary material for "Neurocognitive Heterogeneity in Social Anxiety Disorder: The Role of Self-Referential Processing and Childhood Maltreatment": SM

### Supplementary Information

#### Supplementary Methods

##### Participant screening

Clinical interviews were conducted by doctoral clinical psychologists and doctoral students in clinical psychology trained on the Anxiety Disorders Interview Schedule for DSM-IV: Lifetime version[1]. Patients met criteria if they endorsed greater than moderate social fear in 5 or more distinct social situations assessed by the Anxiety Disorders Interview Schedule for DSM-IV: Lifetime version. Patients also had a score greater than 60 on the Liebowitz Social Anxiety Scale Self-Report, the cutoff score for the generalized subtype of SAD as determined by receiver operator characteristics analysis. Participants were excluded for comorbid diagnoses of current major depressive disorder, posttraumatic stress disorder, or obsessive-compulsive disorder; use of pharmacotherapy or psychotherapy during the past year; participation in cognitive behavioral therapy for any anxiety disorder during the last 2 years; history of neurological disorders or head trauma; cardiovascular disorders, thought disorders, or bipolar disorder; and current substance/alcohol abuse or dependence.

##### Self-Referential Encoding Task

Participants performed a self-referential encoding task programmed in EPrime software (Psychology Software Tools, Pittsburgh, PA). Stimuli were 25 positive and 25 negative social trait adjectives from the Affective Norms for English Words database[2] that were balanced (all ps > 0.51) on word frequency (positive adjectives = 40.5, negative adjectives = 33.6), number of letters (positive adjectives = 6.9, negative adjectives = 7.2), arousal (positive adjectives = 5.54, negative adjectives = 5.43; on a scale of 1 = low to 9 = high), and valence (deviation from neutral: positive adjectives = 2.66, negative adjectives = 2.58; on a scale of 1 = most negative, 5 = neutral, 9 = most positive) based on the nine-point Self-Assessment Manikin[3] rating system. Participants viewed the trait words and made a yes/no judgment indicating whether the trait was self-descriptive ('self' judgment condition), or a yes/no judgment indicating whether the trait was written in all upper-case letters ('case' judgement condition). There were four trial types (self versus case judgment x positive versus negative valence). Each adjective was presented twice, once in the self-referential judgment condition and once in the case judgment condition. The self-referential condition required participants to engage in self-focused attention and access their self-schema in relation to social-evaluative processing. The case identification condition required attention to the concrete sensory features of the words, and was used as a comparison condition to control for reading negative and positive adjectives.

**Supplementary Results**

Table 1. *Descriptive Statistics for Demographic Variables by Samples and Clusters*

| | Samples | | | |
|---|---|---|---|---|
| | Discovery | | Replication | |
| | M | SD | M | SD |
| Age | 33.53 | 8.56 | 32.94 | 8.08 |
| Years of education | 16.81 | 2.13 | 16.83 | 2.77 |
| | N | % | N | % |
| Sex | | | | |
| *Females* | 46 | 48.4 | 51[1] | 53.1 |
| *Males* | 49 | 51.6 | 45 | 46.9 |
| SRET task | | | | |
| *Negative words* | 53.98 | 25.30 | 54.43 | 24.87 |
| *Positive words* | 40.85 | 22.99 | 41.07 | 22.88 |

| | **Clusters discovery sample** | | | | | | | |
|---|---|---|---|---|---|---|---|---|
| | Cluster 2 (positive self) | | | Cluster 1 (negative self) | | | | |
| | M | SD | N | M | SD | N | t | df |
| Age | 34.17 | 9.23 | 42 | 33.02 | 8.05 | 53 | .65 | 93 |
| Years of education | 17.21 | 2.11 | 41 | 16.49 | 2.11 | 50 | 1.63 | 89 |
| | Positive self | | | Negative self | | | | |
| | | | N | | | N | χ2 | *P* value |
| Sex (N) | | | | | | | .02 | .89 |
| *Females* | | | 20 | | | 26 | | |
| *Males* | | | 22 | | | 27 | | |

[1] Sex for one participant was not reported in the replication sample.

| | | | | | | | | |
|---|---|---|---|---|---|---|---|---|
| | **Clusters replication sample (behavior-based clustering)** | | | | | | | |
| | Cluster 2 (positive self) | | | Cluster 1 (negative self) | | | | |
| | M | SD | N | M | SD | N | t | df |
| Age | 34.21 | 8.32 | 25 | 32.49 | 8.00 | 71 | .92 | 94 |
| Years of education | 16.88 | 1.62 | 25 | 16.81 | 3.09 | 70 | .11 | 93 |
| | Positive self | | | Negative self | | | | |
| | | | N | | | N | χ2[2] | *P* value |
| Sex (N) | | | | | | | .11 | .74 |
| *Females* | | | 14 | | | 37 | | |
| *Males* | | | 11 | | | 34 | | |
| | **Clusters replication sample (behavior and brain-based clustering)** | | | | | | | |
| | Cluster 2 (positive self) | | | Cluster 1 (negative self) | | | | |
| | M | SD | N | M | SD | N | t | df |
| Age | 34.08 | 7.86 | 32 | 32.60 | 8.33 | 60 | .83 | 90 |
| Years of education | 17.34 | 2.34 | 32 | 16.53 | 3.04 | 59 | 1.32 | 89 |
| | Positive self | | | Negative self | | | | |
| | | | N | | | N | χ2 | *P* value |
| Sex (N) | | | | | | | 1.02 | .31 |
| *Females* | | | 19 | | | 29 | | |
| *Males* | | | 13 | | | 31 | | |

[2] Chi-square test

Table 2. *Descriptive Statistics and Correlations by Samples*

| | Mean; *SD* (Discovery sample) | 1 | 2 | 3 | 4 | 5 | 6 | 7 | 8 | 9 | 10 | Mean; *SD* (Replication sample) |
|---|---|---|---|---|---|---|---|---|---|---|---|---|
| 1. Emotional abuse and neglect | 11.32; 4.70 | --- | .61*** | .28** | .16 | -.14 | .31** | .06 | -.02 | -.06 | -.16 | 11.74; 4.42 |
| 2. Physical abuse and neglect | 8.40; 2.19 | .56*** | --- | .33*** | .15 | -.06 | .21* | .04 | -.19 | -.12 | -.17 | 7.73; 2.74 |
| 3. Sexual Abuse | 6.23; 3.43 | .19 | .22* | --- | .02 | .07 | -.04 | .11 | -.00 | -.17 | -.31** | 6.02; 3.18 |
| 4. Social anxiety severity | 84.52; 19.33 | .14 | .22* | .18 | --- | -.15 | .24* | -.28** | -.19 | -.01 | -.00 | 90.18; 17.20 |
| 5. Satisfaction with life | 16.85; 7.12 | -.51*** | -.26** | -.19 | -.20 | --- | -.56*** | .22* | .09 | -.11 | .06 | 16.08; 7.56 |
| 6. Stress | 10.05; 3.06 | .24 | .20 | .15 | .37** | -.57*** | --- | -.14 | -.19 | .23* | .01 | 10.46; 3.10 |
| 7. Reappraisal frequency | 3.70; 1.41 | -.11 | .01 | -.02 | .03 | .22* | -.21 | --- | .61*** | -.19 | -.05 | 3.79; 1.27 |
| 8. Reappraisal self-efficacy | 3.38; 1.22 | -.20 | -.05 | -.30 | .27 | .17 | .08 | .71*** | --- | -.22* | .07 | 3.84; 1.41 |
| 9. Suppression frequency | 5.17; 1.33 | .12 | .08 | .09 | .14 | -.23* | .03 | .10 | -.07 | ---- | .59*** | 4.61; 1.28 |
| 10. Suppression self-efficacy | 4.39; 1.55 | -.08 | .11 | .03 | .07 | -.09 | -.01 | .11 | .14 | .52*** | ---- | 4.90; 1.20 |

Note: Discovery sample is represented below the diagonal and replication sample is above

#### Supplementary Figures

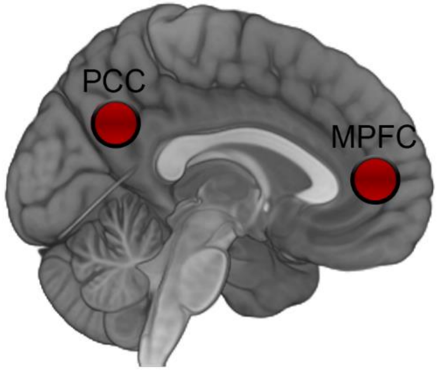


**Figure S1**. Default mode network (DMN) regions of interest (ROIs). For each participant, we averaged mean beta parameter estimates from 10mm spherical ROIs centered on the two hub regions of the DMN: the medial prefrontal cortex (MPFC) and posterior cingulate cortex (PCC). Coordinates were derived from a meta-analysis of self-referential processing (Northoff et al. 2006).

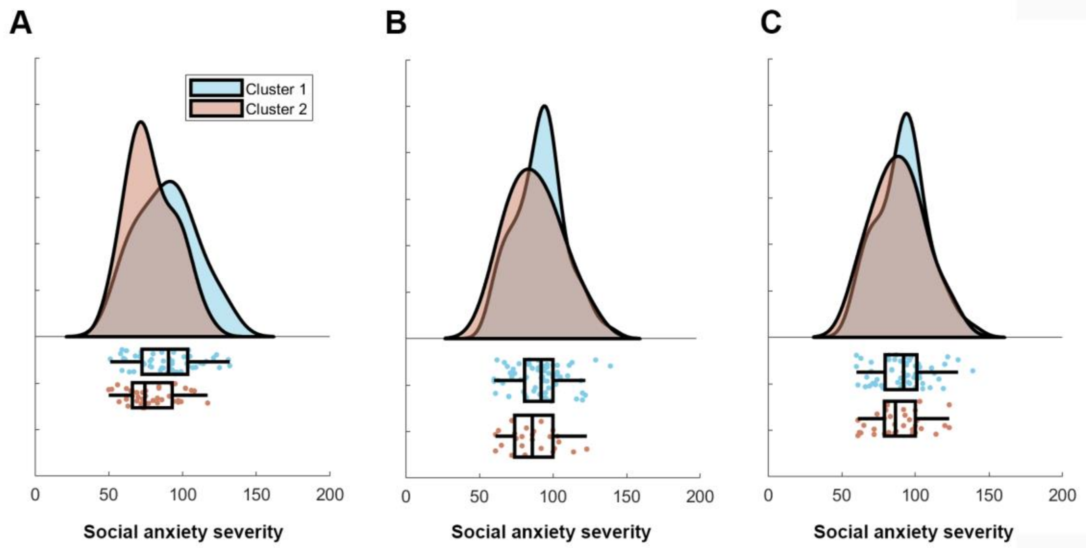


**Figure S2**. Self-reported social anxiety severity (measured with the LSAS-SR) as a function of SAD cluster. (A) Discovery sample. (B) Replication sample (clustering based on behavioral data alone). (C) Replication sample (clustering based on behavioral and brain data).

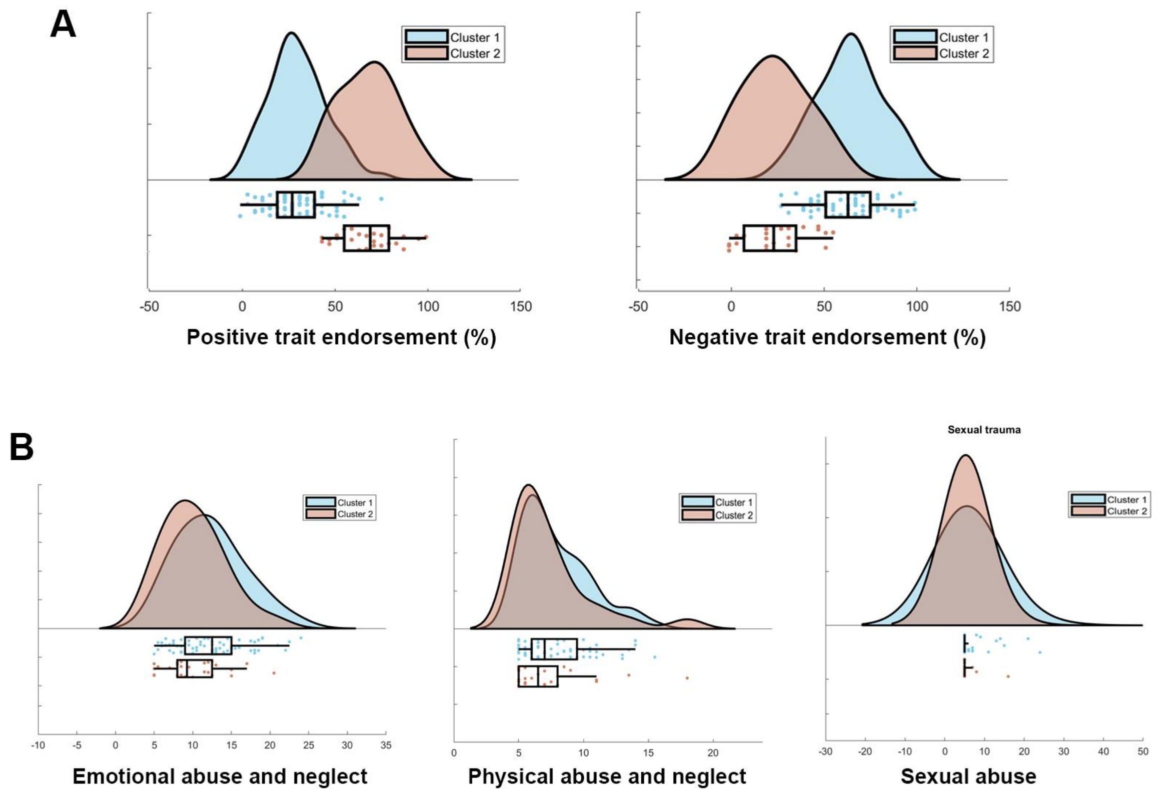


**Figure S3**. Replication sample SAD clusters show distinct profiles of positive and negative self-referential trait endorsement. Clusters were defined based on behavioral performance without brain data (i.e., only based on self-referential trait endorsement). (A) Cluster 1 compared to cluster 2 patients were less likely to endorse positive traits and more likely to endorse negative traits. (B) The clusters also differed in exposure to emotional abuse and neglect, but not physical abuse and neglect or sexual abuse.

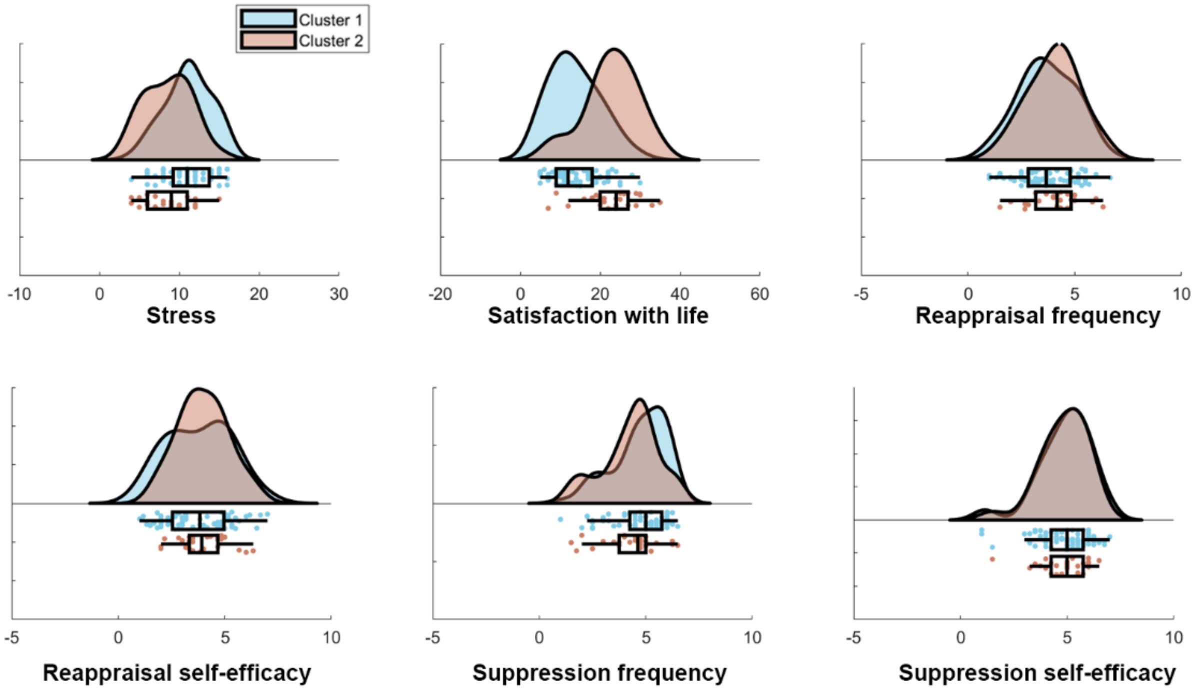


**Figure S4**. Subjective well-being and emotion regulation strategy use as a function of SAD cluster in the replication sample.
